# Impact of ceftiofur administration and *Escherichia coli* inoculation on the calf fecal microbiome

**DOI:** 10.64898/2026.02.09.704862

**Authors:** Andrew J. Sommer, Marta Ferrandis-Vila, Svenja Mamerow, Christian Berens, Christian Menge, Shaodong Wei, Qinqin Wang, Frank M. Aarestrup, Saria Otani, Panagiotis Sapountzis

## Abstract

The cattle gastrointestinal tract harbors a diverse community of microorganisms, including pathogenic and commensal strains of *Escherichia coli*. Antimicrobial use in cattle can disrupt the gut microbiome leading to shifts in bacterial diversity and abundance. Here, we combined shotgun metagenomics and single-cell sequencing to assess how ceftiofur antibiotic treatment impacted microbial diversity and structure. At the start of the experiment, ceftiofur was administered intramuscularly in parallel with the inoculation of a cocktail of extended-beta-lactamase–producing *E. coli* strains, to simulate environmental exposure and acquisition of resistant strains while animals are under antibiotic treatment. Fecal samples were collected from both the antibiotic-treated (ceftiofur and inoculation) and control (inoculation only) calves over the course of 35 days. Read mapping to genome and gene databases showed substantial differences in microbial richness and beta diversity between treatment groups. Treatment group-enriched taxa included *Bacteroidaceae* and *Fibrobacter*, which were more abundant in samples that did not receive ceftiofur, and *Akkermansia* in ceftiofur-treated calves. In ceftiofur-exposed animals, we observed a gradual loss of virulence factors alongside increased abundances of beta-lactam resistance genes, including *cfxA5* and *cfxA6* likely encoded by CAG-485 (*Muribaculaceae*). We further profiled individual cells using single-cell sequencing, which revealed a high number of *Clostridium* carrying macrolide resistance genes *lnu*(P) and *mph*(N) in both ceftiofur-treated and control samples. Overall, our complementary approaches reveal distinct remodeling of the calf microbiome following antibiotic and *E. coli* administration, tied to key functional genes that can be assigned to specific genera or recurrently detected across diverse taxa.

**IMPORTANCE:** Cattle serve as natural reservoirs of zoonotic strains of *E. coli*, which can cause severe gastrointestinal infections in humans. Antibiotic usage on cattle farms can drive the emergence of antimicrobial resistant bacterial strains and alter the underlying cattle gastrointestinal microbiome. Consequently, there is a need to understand how antibiotic administration impacts population dynamics of cattle rumen and intestinal microbes. In this study, we combined both shotgun metagenomics and single-cell genomics on feces from ruminating calves to determine microbiome changes following administration of both ceftiofur and *E. coli* cocktails. We observed considerable variation in prevalence and abundance of virulence factors, antimicrobial resistance–related genes, and taxa with key roles in animal nutrition and health between the microbiomes of antibiotic-treated and antibiotic-free calves, with potential implications for their subsequent development and overall well-being.

## INTRODUCTION

The emergence and spread of zoonotic pathogens and antimicrobial resistance represent a significant threat to global public health (1, 2). While cattle farms are vital for food production, ruminant species may serve as reservoirs for pathogenic strains of *E. coli* responsible for major outbreaks of food-borne illness (3). Livestock farming has also been linked to the spread of antimicrobial resistance genes (ARGs) from commensal bacterial strains to zoonotic pathogens (4, 5), which further necessitates the need to understand interactions between *E. coli* and other resident gut microbes within the cattle gastrointestinal tract.

The cattle gut microbiome can be colonized by phylogenetically diverse lineages including *E. coli* strains, which may encode a range of ARGs and virulence factors (VFs) (6–8). Despite the high genotypic diversity of *E. coli*, disease-causing strains of *E. coli* are broadly classified into distinct pathotypes characterized by specific virulence genes and pathologies. Intestinal pathogenic *E. coli*, including enterohaemorrhagic (EHEC) and other Shiga toxin–producing *E. coli* (STEC), enterotoxigenic *E. coli* (ETEC), enteropathogenic *E. coli* (EPEC) and diffusely-adherent *E. coli* (DAEC), are causative agents of diarrhea in many different mammalian species (9, 10). Differences in *E. coli* pathogenicity across hosts are driven by the ability of strains to attach to and invade host epithelial cells as well as compete with the native microbiota, including commensal *E. coli* strains (11–13). In adult cattle, many intestinal pathogenic *E. coli* asymptomatically colonize the gastrointestinal tract of adult ruminants and are spread to the environment via feces (14). Strains of ETEC, however, can cause severe diarrheal infections in calves and are a leading cause of neonatal mortality on cattle farms (15).

The calf gut microbiome plays an important role in priming the host immune system, the breakdown of nutrients, the development of an anaerobic gut environment, and preventing zoonotic and animal pathogens (such as diverse *E. coli* pathotypes) from colonizing the intestinal environment via microbial competition (14, 16, 17). Perturbations to the gut microbiome during development, including bacterial infections and subsequent administration of antimicrobials, can lead to shifts in microbial communities and a decrease in overall bacterial diversity (18, 19). However, despite prior investigations, it remains unclear how these changes can affect the intestinal microbiota in the strain- and species-level, during and post treatment, and how the genomic inventory (including key-genes related to antimicrobial resistance and virulence potential) of the microbiome is impacted. Thus, longitudinal genomic profiling of the cattle intestinal microbial community can provide valuable insights into how specific lineages adapt, compete, and colonize the gut under shifting environmental conditions.

Within this framework, the HECTOR (*Host restriction of Escherichia coli on Transmission dynamics and spread of antimicrobial Resistance*) consortium performed large-scale whole genome sequencing of *E. coli* isolates to identify potential colonization determinants or host associated strains across different mammalian species (20). While this work has identified specific gene clusters associated with different animal hosts (20), this has not always been the case (21). Additionally, because ESBL–producing *E. coli* are of particular concern in both veterinary and human health, and because ceftiofur—a third-generation cephalosporin with broad-spectrum activity—is widely used in the cattle industry to treat metritis, mastitis, and bovine respiratory and diarrheal diseases (49, 50), we examined microbial changes on young calves under ceftiofur exposure and in the presence of ESBL strains, which simulated the introduction of clinically relevant resistance determinants under controlled conditions, thereby providing a standardized background against which antibiotic-driven microbiome and resistome shifts could be assessed.

Following the administration of ceftiofur and ESBL *E. coli* strain on calves, we performed shotgun metagenomic and single-cell sequencing (SCS) on their fecal samples to investigate how ceftiofur administration alters gut microbiome communities. We further characterized both the virulome (identified virulence genes) and resistome (identified ARGs) to determine how temporal changes and antimicrobial resistance can lead to shifts in clinically relevant genes in the calf gastrointestinal tract.

## METHODS

### Animal rearing

Animal husbandry and experimental procedures were conducted in accordance with protocols approved by the State Office for Agriculture, Food Safety and Fisheries of Mecklenburg-Western Pomerania, Rostock, Germany (reference no. 7221.3-1-103019/20-1). A total of twenty *Holstein Friesian* ruminating calves aged between 10-18 weeks-old were housed in an environmentally controlled biosafety level 2 animal facility at the Friedrich-Loeffler-Institut (FLI) on the Isle of Riems, Greifswald. Upon arrival, all animals tested negative for ceftiofur-resistant *Enterobacteriaceae* by plating fecal samples on ceftiofur-containing (4 μg/ml) Gassner*-agar* plates.

Two sets of animal experiments were conducted from *timeframe1* (NJ) and *timeframe2* (GWAS). For each experiment, ten calves were divided into two treatment groups (n=5 animals each) and housed separately with one group receiving supplementary antibiotic treatment and the other serving as an untreated control. Both groups were fed a need-based diet of hay cobs and concentrated feed. Following a three-week acclimation period, the antibiotic treatment group received daily intramuscular injections of ceftiofur (cephalosporin: 1 mg/kg body weight; EXCENEL Flow®, Zoetis, Parsippany, NJ, USA) for three consecutive days (Experimental day - 01, 00, +01) while all control group animals were administered intramuscular injections of sterile saline solution (2ml, irrespective of their body weights). One calf in the GWAS ceftiofur-treated group died shortly after inoculation, reducing the number of animals in that group from ten to nine, while the NJ group number of animals (n=10) remained unchanged. Experimental inoculation of the *E. coli* strains was performed on all animals intra-ruminally via nasogastric tube on experimental day 00. Briefly, cocktails were prepared by combining exponential phase liquid cultures of selected *E. coli* strains and combined in equal proportions, resulting in a final inoculant of approximately 1 x 10^10^ bacteria. The combined inoculants were centrifuged to pellet bacterial cells before resuspension in a 10% sodium bicarbonate saline solution. Details on the cocktail strain metadata are available as supplemental material (Table S1). Fecal samples (∼50 mL) were collected in sterile Falcon tubes via rectal sampling over the course of both experiments (Table S2), with the initial sample collected immediately prior to ceftiofur administration on day -01. All samples were stored at −80 °C until further processing.

### Shotgun metagenomic sequencing and Metagenome Assembled Genome (MAG) reconstruction

Genomic DNA was extracted from the 77 fecal samples using the QIAamp Fast DNA stool mini kit (Qiagen, Germany) following the manufacturer’s instructions. From each microbiome, 200 mg feces was used and DNA was eluted in 50 µL of pre-heated (65°C) AE buffer to increase DNA yield. DNA quality was checked using a Qubit Fluorometer (Thermo Fisher Scientific). For metagenomics sequencing, all libraries were prepared using the PCR-free Kapa Hyper Prep Kit (Roche). All libraries were sequenced on the Illumina Novaseq 6000 S4 (2 × 150 bp) platform with an estimated output of 46 million reads per sample (average).

Retrieved sequences were filtered to remove low quality reads and adaptor sequences using Trimmomatic before further analysis (22). Trimmed reads were assembled into contigs with a minimum length of 300 bp using MEGAHIT (23). Contigs that were at least 1500bp or longer were binned into MAGs using MetaBAT (24). From the generated bins, we retained only high quality bins with >90% completeness and <5% contamination, as assessed by CheckM (25). MAGs were dereplicated using dRep with primary and secondary ANI thresholds of 0.97 and 0.99 to generate a non-redundant MAG dataset (26). The specificity of this database was determined by evaluating the percentage of reads aligned to the MAG dataset via a global alignment approach (26) and a kmer signature approach (27), using default settings. We additionally identified the Open Reading Frames of all reconstructed MAGs using prodigal (28) and the predicted amino acid sequences were used as input to: i) perform diamond blast comparisons (29) against the ResFinder (obtained October 2022) and the Virulence Factor (VFDB) databases (obtained July 2023) (30, 31).

### Analysis of microbial community structure

The dereplicated MAG dataset (n=1607 MAGs) was used as a reference database for KMA (K-mer alignment) to create a count table of estimated MAG abundance and coverage across all NJ group calf samples (32). Count tables were normalized by creating ratios using the gene length and genome counts calculated with MicrobeCensus (33). The resulting abundance count table was imported into R (version 4.4.2) and filtered to exclude hits with less than 10% genome coverage. Observed MAG richness was determined by counting the number of unique MAGs found across each sample, and Shannon diversity was estimated using the ‘vegan’ (version 2.6.10) package in R (34). Statistical differences in alpha diversity between treatment groups at each sampling point were assessed using a Wilcoxon rank-sum test with Bonferroni correction for multiple comparisons.

Bacterial abundance counts were imputed to remove samples or MAGs with less than 20% prevalence using ‘zComposition’ in R (version 1.5.0-4, method “CZM”) and transformed to center log-ratios (CLR) (35). Sample dissimilarity was calculated using Euclidean distances of CLR-transformed MAG abundances with the ‘vegan::vegdist’ function and visualized through Principal Coordinates Analysis (PCoA). Statistical differences in bacterial beta-diversity between groups were determined through permutational multivariate analysis of variance (PERMANOVA) using the ‘vegan::adonis2*’* function. Differences in dispersal between tested groups were determined via permutational multivariate analysis of dispersion (PERMDISP) using the ‘vegan::betadisper’ function (34). To determine MAGs that were enriched in either ceftiofur-treated or control samples post-antibiotic administration, we performed differential abundance tests on normalized counts of MAG abundance counts using the ‘indicspecies’ package in R (version 1.8.0, 999 permutations) (36). All p-values were adjusted using a false discovery rate (fdr) correction. Only MAGs with an IndVal stat greater than 0.80 and a fdr adjusted p-value less than 0.05 were considered highly enriched to a specific treatment group. Visualization of abundance and diversity metrics was performed in R using the ggplot2 package (version 3.5.1) (37).

### Analysis of resistome and virulome

Identification of VFs and ARGs were performed by aligning the metagenomic reads using KMA against reference databases constructed from nucleotide sequences obtained from VFDB and ResFinder (30–32). Count tables were then normalized and transformed using MicrobeCensus and imported into R for analysis (33). Count data were first transformed to remove all hits with less than 90% coverage to a reference gene, and subsequently imputed so that samples or genes present in less than 15% of the cases were excluded. The imputed count tables were subsequently CLR transformed as described above. Statistical differences in ARG abundance between treatment groups were assessed using a Wilcoxon rank-sum test with Bonferroni correction for multiple comparisons. No CLR transformation or statistical test was performed on VFs as most identified genes were rare.

### Single-cell Sequencing

Due to its computationally demanding nature, single-cell sequencing was performed on a subset of the calves where longitudinal sampling allowed us to follow treatment over time. The sequencing was performed using the semi-permeable capsule (SPC) workflow as described previously (38) with minor adaptations to bovine fecal samples. Briefly, bacterial cells were detached from fecal material, filtered, and encapsulated into SPCs using the ONYX microfluidics platform (Atrandi Biosciences) targeting a λ value of 0.1 to minimize multiple occupancy. Encapsulated cells were subjected to sequential enzymatic and proteinase lysis, followed by multiple displacement amplification (MDA) within the SPCs. DNA was then combinatorially barcoded through a split-and-pool strategy, using 16 × 96 × 96 × 96 barcode combinations, enabling the generation of thousands of single-amplified genomes (SAGs) per run. After release from SPCs, barcoded DNA was purified, prepared into Illumina-compatible libraries (NEBNext Ultra II FS kit), and sequenced on an Illumina Novaseq 6000 S4 platform. Sequencing data were demultiplexed into individual SAGs using high-dimensional barcode deconvolution, trimmed with fastp (39) similar to Ling et al. (38). Identification of ARGs were performed by aligning each SAG paired reads against the ResFinder database using KMA, as described above (31, 32). Only hits with at least 90% coverage against the reference template were retained for further analysis. Taxonomic identification of SAGs was performed using Sourmash against the GTDB database (27, 40).

## RESULTS

### Construction and validation of a bacterial MAG database

Shotgun metagenomic sequencing was initially performed on the total of 77 calf fecal samples (Table S2; NJ and GWAS experiments), generating an average of 46 million reads per sample (range 39-68 million reads). Sequence reads were assembled into a total of 1,607 MAGs with a filtering threshold of >90% completeness and <5% contamination (Table S3A). Genus-level identification was assigned to 1,219 MAGs, representing 128 bacterial genera and one archaeal genus. The most prevalent bacterial genera in the MAG database included *Alistipes* (n=238), *Cryptobacteroides* (n=81), *Akkermansia* (n=58), and *Treponema*_D (n=52); 368 MAGs could not be assigned to bacterial taxa. Mapping of reads to the 1,607 reconstructed MAGs showed an average alignment rate of 43.4% (median=43.45%; range=33.9-51.5) across all samples, suggesting that the reconstructed MAGs covered nearly half of all sample reads. A weighted-abundance approach of mapping Kmer signatures showed that almost 35.7% of reads could be mapped to our MAG database compared to 22.9% of reads against Genbank sequences (medians), confirming the improved specificity of the custom MAG database.

### Variation between control and ceftiofur-exposed calf microbiomes

During the administration experiment, the GWAS animals did not complete the full sampling timeline (after ceftiofur administration, samples were obtained only at day 3 and day 14 instead of through day 35). Additionally, preliminary mapping of the GWAS sample reads to the MAG database revealed substantial baseline differences in microbiome composition between treatment groups (ceftiofur and control), limiting the interpretability of treatment-associated changes (Figure S1A). For these reasons, the GWAS group was excluded from all further analyses, and we focused on the total of the 50 metagenome samples collected from 10 animals (NJ experiment) reared over the 35-day period (Table S2).

Reads from both the ceftiofur-treated (n=25 samples: 5 animals x 5 time points) and control groups (n=25 samples: 5 animals x 5 time points) were mapped to our custom MAG database to assess the impact of antibiotic exposure on bacterial communities and MAG abundance. Analysis of bacterial alpha diversity revealed a marginally significant difference in MAG richness between control and ceftiofur-treated calves on day five (Bonferroni p=0.0397), which corresponded to the first sample collected following cessation of antibiotic administration (Figure 1A). No significant differences in alpha diversity using Shannon Index were observed between treatments at any timepoint, even though a non-significant small difference between control and ceftiofur samples was present again.

**Figure 1.**
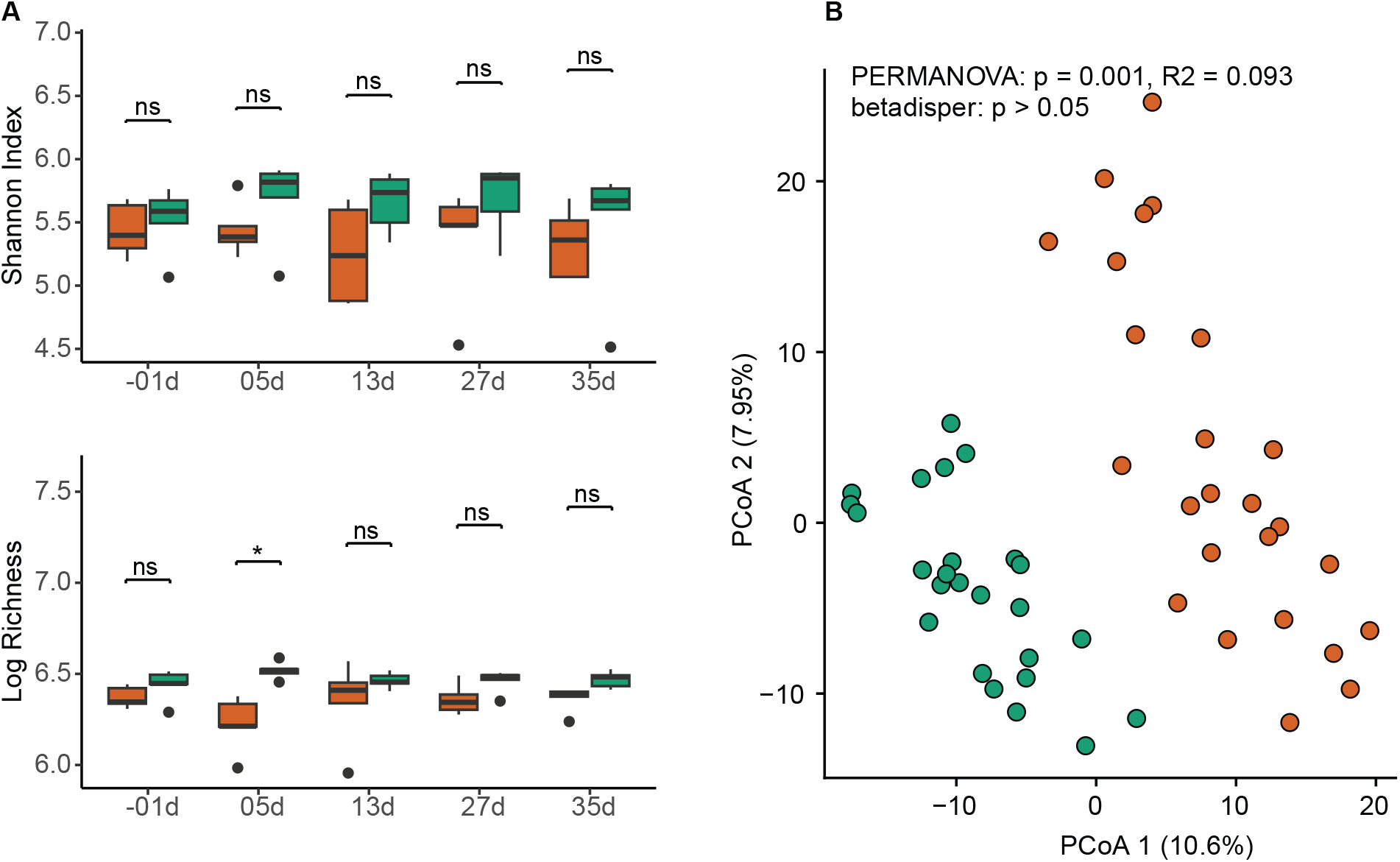
MAG level alpha- and beta-diversity analysis of calf fecal samples. (**A**) Boxplots showing Shannon diversity index and log-transformed richness of MAGs over time. Statistically significant differences in alpha-diversity metrics between the control and ceftiofur-treated groups at each timepoint were determined via Wilcoxon rank-sum test (p<0.05, Bonferroni correction). (**B)** PCoA of beta diversity based on Euclidean distances of MAG abundance counts. Differences between groups were assessed using PERMANOVA, and group dispersion was evaluated using betadisper. Ceftiofur samples are shown in orange; control samples are shown in green.

Principal coordinates analysis (PCoA) of beta diversity based on Euclidean distances of MAG abundance counts revealed distinct clustering of the bacterial communities because of the antibiotic treatment (Figure 1B). We found that antibiotic-exposed and control microbiomes differed significantly in centroid positions (PERMANOVA, *p*<0.05), while displaying no significant differences in within-group dispersions (betadisper, *p*>0.05). While similar patterns were also observed for both sampling date and calf ID number (PERMANOVA, *p*<0.05; betadisper, *p*>0.05), visual clustering was less apparent (Figure S1C).

To identify MAGs enriched between antibiotic-exposed and control microbiomes, we performed a differential abundance test (indicspecies). We identified a total of 49 MAGs highly enriched in the control group and 17 MAGs highly enriched in the ceftiofur group (fdr<0.05; stat>0.80) (Table S3B). The ceftiofur-associated MAGs belonged to the following genera: *Akkermansia* (n=7), *Alistipes* (n=1), *Barnesiella* (n=3), *Lachnospiraceae* CAG-791 (n=2), *Paludibacteraceae* RF16 (n=4) (Figure 2A), while MAGs enriched in the control group belonged to 23 distinct genera from 17 distinct bacterial families (Figure 2B): *Limimorpha* (n=10), *Bacteroidaceae* HGM04593 (n=6), *Paludibacteraceae* UBA4363 (n=5), and *Cryptobacteroides* (n=4), along with several other taxa, notably including *Fibrobacter* (n=1), *Ruminococcus* (n=1), and *Alistipes* (n=1).

**Figure 2.**
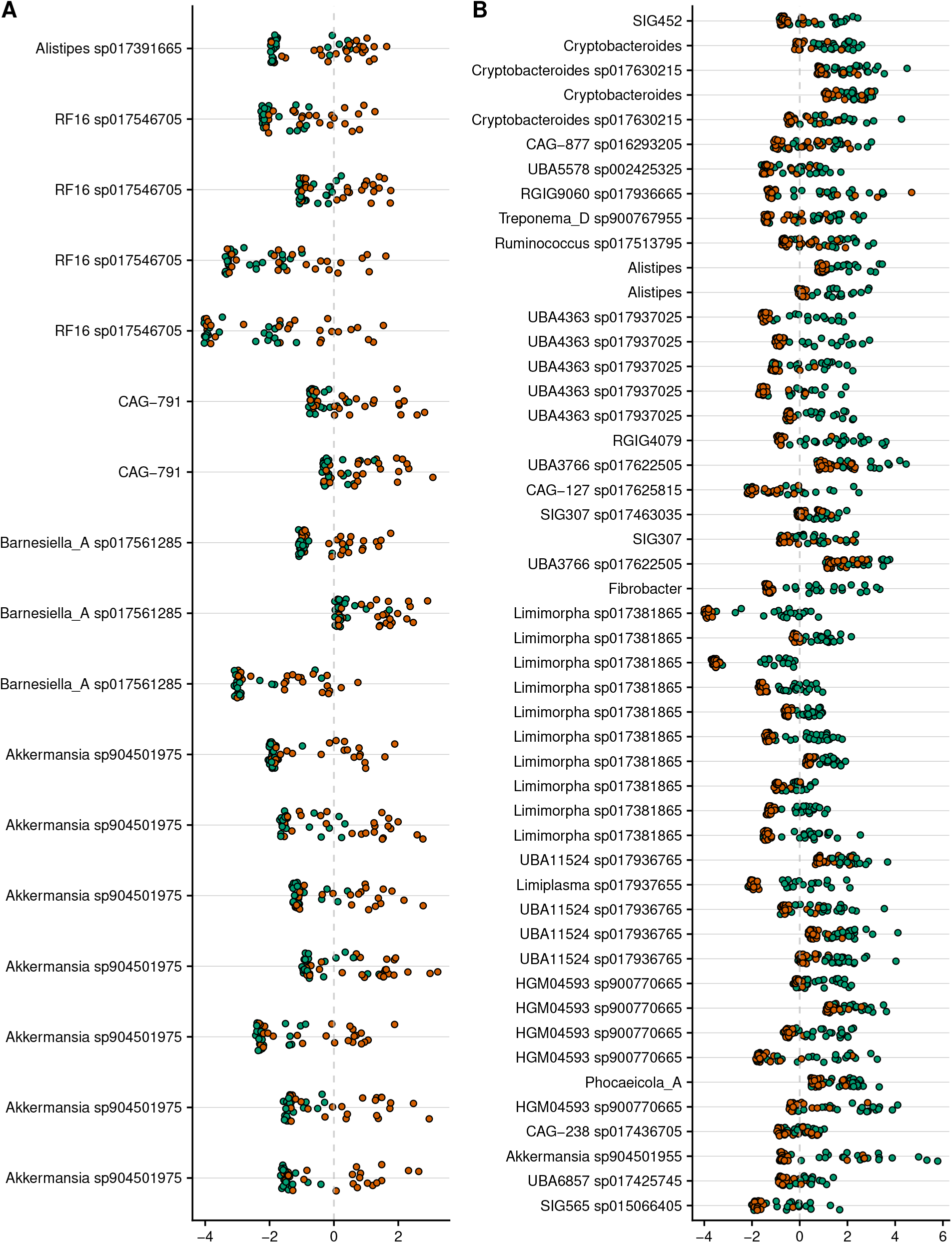
Identification of MAGs associated with either ceftiofur-exposed (**A**) or control (**B**) fecal samples from calves. The CLR transformed abundances of each MAG in individual ceftiofur-exposed (orange) and control (green) samples are shown for MAGs classified as highly enriched to either group (IndVal>0.80; fdr adjusted p<0.05).

We further examined the most prevalent MAGs within each group to identify core taxa in bovine fecal microbiomes. We found that a total of 130 MAGs were shared between 80% of both ceftiofur-treated and control samples (n=20 per group), with an additional 148 MAGs found in 80% of all control samples and 64 MAGs found in 80% of the ceftiofur-treated samples (Figure S2). The most common bacterial taxa represented by these shared MAGs included *Rikenellaceae* (*Alistipes*), *Bacteroidaceae* (UBA1189, *Phocaeicola, Paraprevotella*, HGM04593), and UBA932 (*Cryptobacteroides, Egerieousia*). None of the 130 highly prevalent shared MAGs were identified as differentially abundant (fdr<0.05; stat>0.80) in either group. We found that many of the 64 MAGs from the ceftiofur-treated group were still found in the control group at a lower prevalence (range 5-19 samples). By contrast, ten of the highly prevalent control group MAGs (8 *Limimorpha*, 1 *Fibrobacter*, 1 Clostridia) were completely absent from the ceftiofur-treated samples (Figure S2).

We further attempted to take advantage of the availability of the administered *E. coli* genome sequences (Table S1) by using the denovo assembled contigs (which were also used for the MAG reconstructon) to map directly to these reference genomes (Supplementary File 1). However, although local BLAST comparisons suggested that genomic fragments of the administered strains may have been present in several samples post administration, a small number of spurious hits identified in samples before the *E. coli* inoculation, and the positive identification of contigs from all reference genomes, suggested that the results are likely biased by highly conserved regions shared between native and administered *E. coli* strains (Supplementary File 1).

### Analysis of virulome and resistome between samples

We next characterized antibiotic resistance genes across samples. We identified a total of 57 unique antimicrobial resistance genes across all samples (gene coverage >90%), which notably included the extended-spectrum beta-lactamase genes *bla*_CTX-M-1_ (n=1 sample) and *bla*_TEM-116_ (n=1 sample). CLR abundance values were calculated for the 27 genes with an overall prevalence of at least 15%, which included 3 aminoglycoside, 5 beta-lactam, 4 macrolide, 2 phenicol, and 12 tetracycline resistance genes (Figure 3A). Abundances of *cfxA5, cfxA6*, and *mph*(N) were significantly higher in the ceftiofur-exposed microbiomes, while abundances of the *cfr*(C) gene were significantly higher in control samples (Wilcox rank sum test, p<0.05, Bonferroni correction). Visualization of gene abundance by sampling date showed a strong rise in the abundance of the *cfxA5* gene in ceftiofur-treated samples post antibiotic administration whereas the abundance of the gene remained relatively stable in control samples (Figure 3B). We next attempted to trace the origin of ARGs across assembled MAGs, which showed that *cfxA5* and *cfxA6* were identified in MAGs classified as CAG-485 (family *Muribaculaceae*; Figure S3). While we did not identify any statistical differences in abundance by treatment in these MAGs (indicspecies, fdr>0.05), the maximum abundance of both CAG-485 MAGs was highest in ceftiofur-treated samples (Figure S4). This difference was most apparent for the CAG-485 MAG encoding *cfxA5*, which showed a large increase in abundance in several samples after day 05.

**Figure 3.**
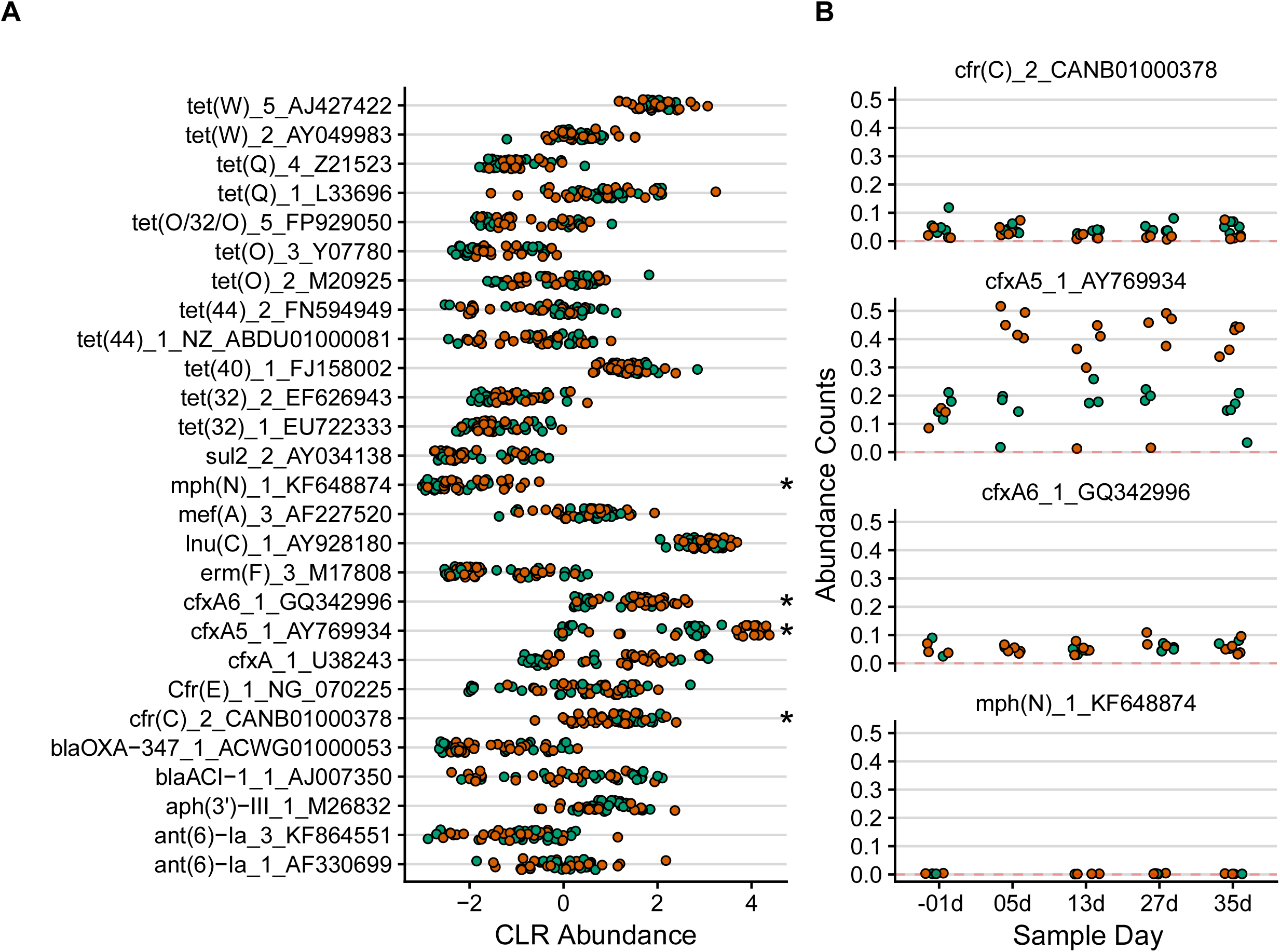
Identification of ARGs across calf fecal metagenomes. (**A**) CLR transformed abundances of all ARGs found in at least 15% of both ceftiofur-exposed (orange) and control (green) fecal samples with over 90% reference gene coverage. Asterisks indicate comparisons with a significantly different abundance between ceftiofur-exposed and control samples (*p* < 0.05, Wilcoxon signed rank test, Bonferroni correction). (**B**) Plots show normalized ARG abundances over time for statistically significant comparisons.

We then performed a similar analysis to quantify bacterial virulence factor abundance through mapping metagenome reads against VFDB (30). While virulence factors meeting the 90% coverage threshold were completely absent from 31 samples, over 100 VFs were identified in three separate control samples (Figure 4A). In total, we identified 247 VFs across all samples, with 191 genes unique to samples from the control group and no genes unique to samples from the ceftiofur group (Figure 4B). The genes identified were grouped by shared function, revealing 24 *Escherichia-Shigella* VF groups present in at least five individual samples (Figure 4C). The most prevalent virulence factor type detected were adhesion factors, which notably included afa/Dr genes, F9 fimbriae, HCP, type 1 fimbriae, and Stg fimbriae. Virulence gene prevalence was highest in the control group during the start of the experiment and declined over time, with no genes detected in any of the samples (control or ceftiofur treated) on day 35 (Figure 4C).

**Figure 4.**
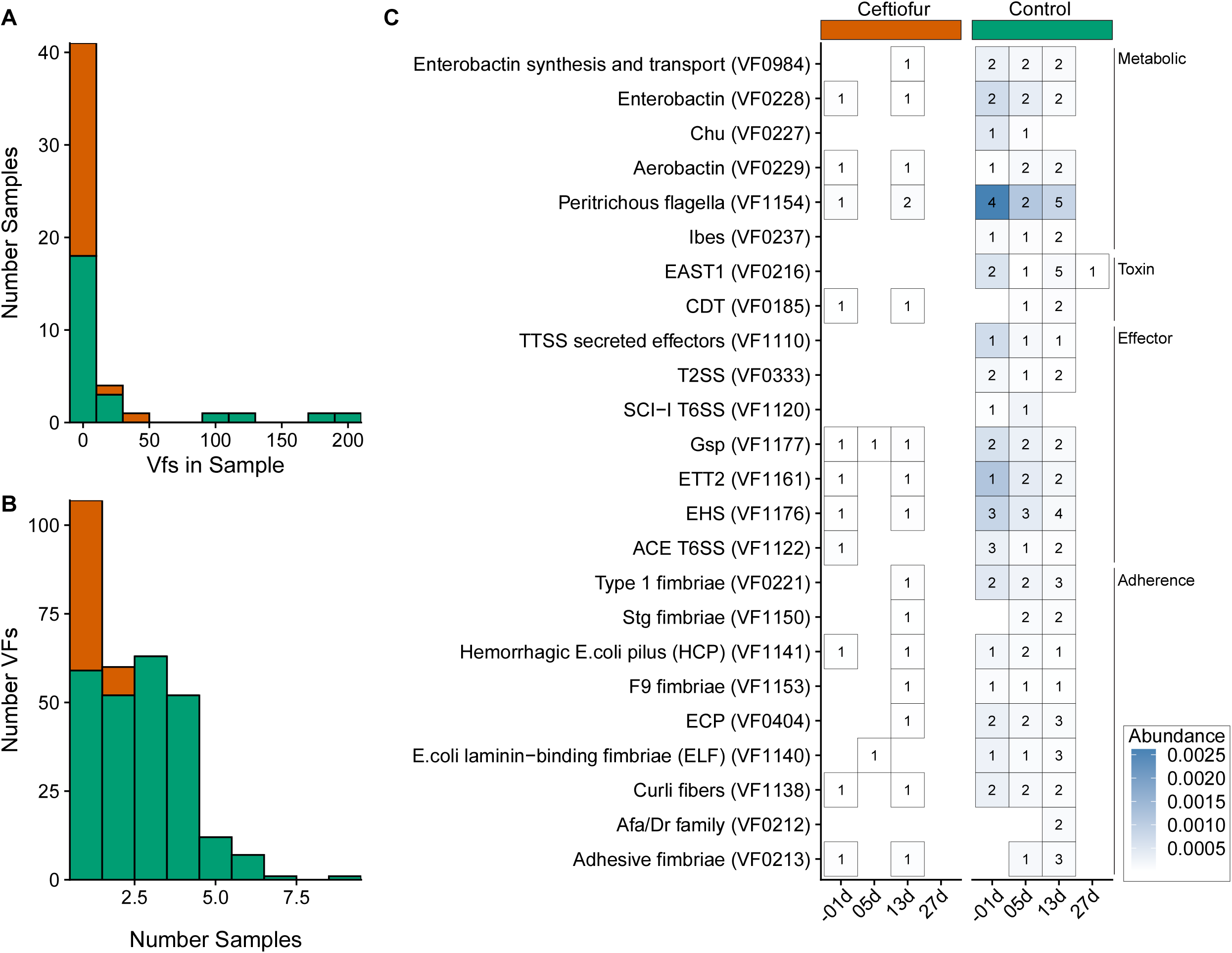
Identification of VFs in calf fecal samples (>90% reference gene coverage). (**A**) Histogram showing the number of virulence factors (VFs) in each fecal sample. (**B**) Histogram summarizing the prevalence of unique virulence factors across fecal samples. (**C**) Heatmap showing the prevalence and abundance of grouped virulence factors (*pathogene*) across sampling day and treatment type. Only VF groups found in at least 5 samples are shown. Numbers inside boxes indicate the number of distinct samples in which the virulence factor was detected at the given time point and treatment.

### Single-cell Sequencing of Bacteria

In parallel with the metagenomic analyses and community characterization based on MAG reconstruction and read mapping, we applied a single-cell sequencing approach to assess individual variability at strain-level resolution. A subset of samples was analyzed to evaluate whether single-cell and metagenomic approaches can yield congruent results. We performed SCS in the feces of two ceftiofur-treated calves and one control calf. Taxonomic classification was obtained for a total of 6,552 out of 10,122 sequenced cells (Table S4). The most common taxonomy of the sequenced cells included *Clostridium* (n=3671 cells), *Romboutsia* (n=906 cells), *Turicibacter* (n=496 cells), unclassified *Peptostreptococcaceae* (n=465 cells), and *Paraclostridium* (n=240 cells) (Figure S5; Table S4). A total of 19 different ARGs were identified across all sequenced cells, with *tetB*(P) (n=585), *mph*(N) (n=547), and *lnu*(P) (n=469) being the most prevalent genes (Figure 5A). The macrolide resistance genes *mph*(N) and *lnu*(P) were primarily encoded by *Clostridium* (Figure 5B). The most common carriers of *tetB*(P) include *Romboutsia* and other unidentified *Peptostreptococcaceae*; however, a portion of cells (35.04%) encoding this gene could not be taxonomically classified (Figure 5B). Other less commonly identified ARGs, including various aminoglycosides, macrolide, and tetracycline resistance genes, were also encoded by cells that could not be taxonomically classified.

**Figure 5.**
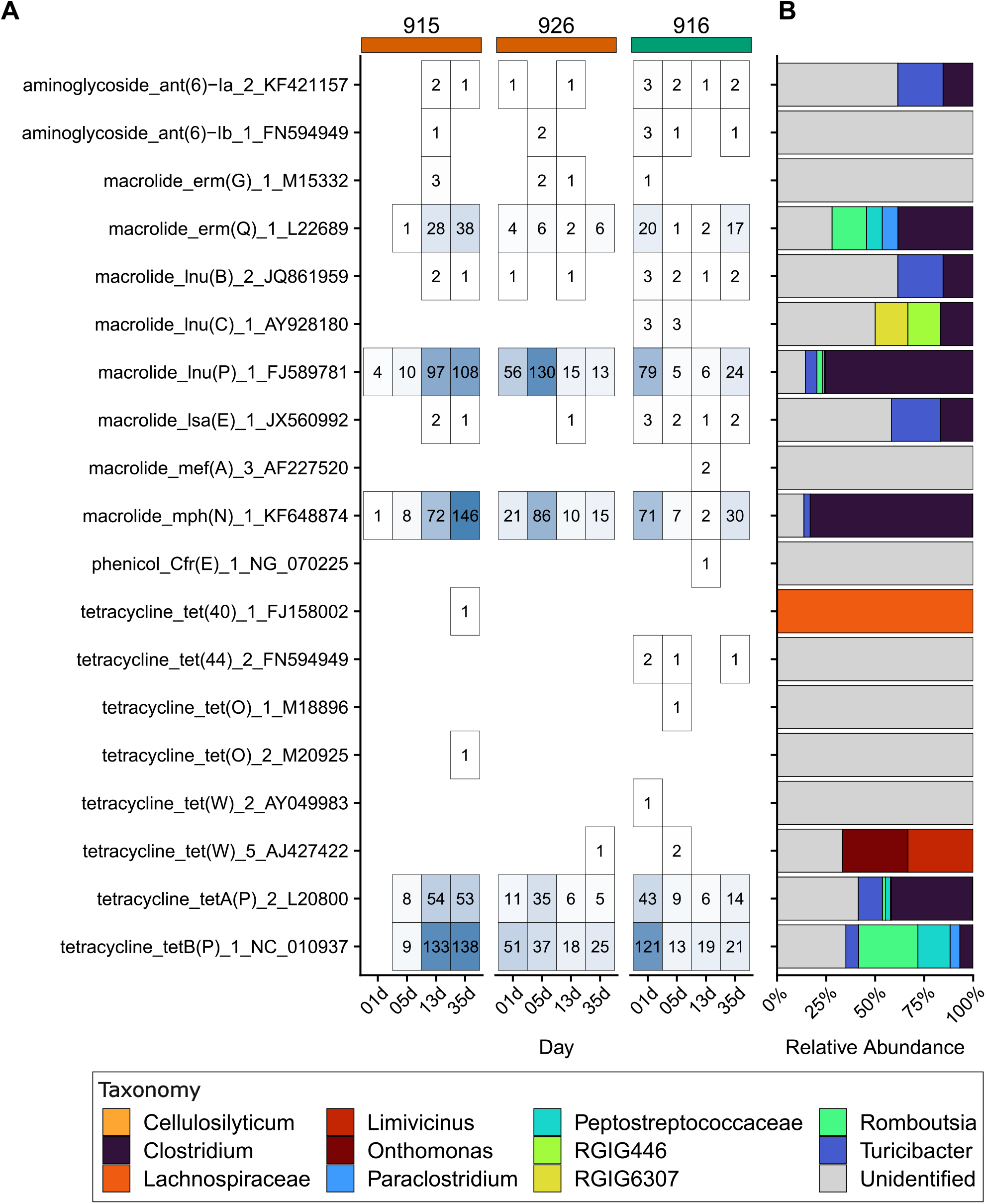
Identification of ARG carriage through single-cell sequencing. (A) Distribution of ARGs identified in fecal samples selected for single-cell sequencing. The number inside each box represents the count of cells encoding the respective ARG with at least 90% coverage against the reference template (B) Taxonomic distribution of bacterial cells encoding each ARG. Bar height corresponds to the relative abundance of bacterial families, or genera when available. Cells not classified to at least the family level are labelled as unidentified.

## DISCUSSION

Here, we applied metagenomic sequencing supplemented with SCS to investigate shifts in the overall microbial community diversity in calf feces post antimicrobial administration. All calves in the experiment were inoculated with a cocktail of genotyped, ESBL *E. coli* strains, which allowed us to examine how their introduction affected the bovine gastrointestinal tract, with and without microbiota perturbation by ceftiofur. Our analysis revealed significant differences between the bacterial communities associated with antibiotic-treated calves compared to control samples. We tried to evaluate the persistence of the administered *E. coli* cocktail using different approaches, but the strain-specific signals did not align sufficiently across methods to support reliable interpretation. More specifically, comparisons between *E. coli* genomes and assembled contigs suggested their presence post inoculation but the highly conserved genomic similarity between *E. coli* strains was likely the main culprit for obscuring precise strain-level identification (Supplementary File 1). SCS on the other hand which could have shown some better insight was biased by the large abscence of gram negative bacteria including *E. coli* (see also below). Therefore, we focused our interpretation on microbiome and resistome dynamics where the data provide consistent and reproducible patterns.

We first assessed the temporal changes in the calves’ fecal microbiome after a three-day course of injectable ceftiofur hydrochloride, a third-generation beta-lactam widely used in buiatrics (41, 42). We observed a significant difference in overall microbial diversity between treatment groups, including a marked decrease in bacterial richness following ceftiofur injections. Although overall bacterial richness remained lower in the ceftiofur-treated samples, no significant differences were observed at later time points, suggesting that the bacterial communities recovered over time. Our findings are largely consistent with previous studies, which observed distinct changes in bacterial alpha diversity during ceftiofur administration, although these studies did not investigate strain-level dynamics in comparable depth (43–45). Further beta-diversity analysis showed clustering of samples by treatment groups during the entire course of sampling, with divergence between groups increasing as the experiment progressed. Despite some pre-existing differences in bacterial beta-diversity, the antimicrobial treatment was still linked to substantial shifts in microbial diversity over time. We identified significant changes in MAG-level abundance between treatment groups post-ceftiofur administration, suggesting that antibiotics could have amplified potential underlying group differences. A greater number of bacterial MAGs were enriched within control samples. Separate MAGs were identified as enriched in control and ceftiofur samples for *Akkermansia* and *Alistipes*, which represent highly diverse bacterial taxa commonly found in the ruminant gastrointestinal tract (46, 47). In our samples we reconstructed a total of 58 *Akkermansia* MAGs and 238 *Alistipes* MAGs, and differences in the abundances between treatment groups likely represents strain-level variations in colonization. In contrast, we identified that control samples were highly enriched in a *Fibrobacter* MAG, a genus which is important for cellulose digestion in the cattle rumen (48), and the observed changes may reflect delayed development of the calf rumen microbiome or altered functional capacity of the gut microbiome. Notably, *Fibrobacter* was absent not only in ceftiofur-treated calves but also from samples collected prior to antibiotic administration, suggesting that this disparity is inherent to the animal groups rather than treatment-induced. Nevertheless, given that only a subset of calves from the control group carried this MAG—and assuming this pattern is not attributable to poor sequencing depth— ceftiofur may have impeded *Fibrobacter* colonization in a group of animals where this taxon was already underrepresented.

The beta-lactamase genes *cfxA5* and *cfxA6* were enriched in ceftiofur samples, while bacteria encoding the *cfr(C)* were enriched in control samples. The *cfr* gene, a methyltransferase often located in various mobile genetic elements, is of growing concern since it provides multi-resistance to various taxa and has been identified in many countries and hosts (49). In our experiment, we assume that the taxa carrying it were either eliminated by the ceftiofur administration (most likely scenario) or were not present in the ceftiofur exposed group. The enrichment of *cfxA5* and *cfxA6* in the antibiotic group may have been similarly driven by shifts in the underlying microbial community structure caused by the antibiotic perturbation. We were able to map these genes to CAG-485 (family *Muribaculaceae*) MAGs, which showed a marked increase in abundance in a subset of ceftiofur-treated samples, although it is also possible that these genes were encoded by other bacteria not represented in our MAG database. Both *cfxA5* and *cfxA6* have been previously reported in other Bacteroidales bacteria, including *Prevotella* and *Bacteroides*, and are associated with resistance to beta-lactam antibiotics (50–52). Prior studies have also reported increases in *cfxA* gene abundance during ceftiofur administration in cattle (43, 44), suggesting that this effect is not unique to our samples. However, other studies have also reported contrasting results, including no changes in beta-lactam resistance during ceftiofur exposure (53). These discrepancies may be partly explained by differences in study design including animal age or breed (54, 55), management practices (56), ceftiofur drug formulation (57), or sample collection methods – all of which collectively impact the fecal microbiome structure.

Finally, considering the observed enrichment of beta-lactamase genes *cfxA5* and *cfxA6* in Muribaculaceae MAGs, while classical ESBL genes such as blaCTX-M-1 and blaTEM-116 were only rarely detected, we tentatively suggest that ceftiofur administration may select for non-Enterobacteriaceae beta-lactamase reservoirs. Unfortunately, the resolution of our data cannot further determine whether this enrichment is part of a shift in the reservoir of beta-lactam resistance from enteric pathogens to commensal taxa, or a broader increase to both populations.

We performed SCS on a subset of samples to detect the carriage of ARGs by individual live bacterial cells. While SCS is a powerful tool that allows for the direct linkage of genes to individual bacterial cells, the profiled cells represent a subset of the total microbial community that survived post-collection storage and were successfully sorted and amplified (38, 58). The lack of sequenced Gram-negative bacterial cells, which likely also explains the absence of beta-lactamases genes in the SCS results, has been previously reported in previous SCS from fecal samples (38) and is therefore likely reflective of a corresponding reduction of cell viability due to freezing of sample material. The most abundant bacterial taxa identified included *Clostridium* and *Peptostreptococcaceae* (including *Romboutsia*), which are common Gram-positive members of the cattle fecal microbiome (47, 59). Our SCS revealed that the *Clostridium* cells encoded several macrolide resistance genes, including *mph*(N) and *lnu*(P), with the former being significantly more abundant in ceftiofur-exposed compared to control microbiomes. However, since none of the MAGs in the database encoded this gene with high coverage, which is likely due to the amplification step employed in the SCS protocol that possibly increased their presence in the single-cell sequencing output. Likewise, we identified that *tetW* (AJ427422) was only encoded by a single *Onthomonas* and a single *Limivicinus* cell. While *tetW* was highly abundant in the metagenomic sequencing data, the gene was not encoded with high coverage in any MAG. This highlights a known limitation of the MAG method, as genomes with high complexity, increased strain variation, or low abundance are more difficult to reconstruct. The increased ability to link genes to individual bacterial hosts highlights the value of pairing metagenomic data with SCS for bacterial community analysis.

Screening of entire metagenomes against virulence factor reference sets revealed many classical pathotype-associated genes were rare across samples, however, we still identified several adhesion factors linked to specific pathotypes (including afa/Dr, Hemorrhagic Coli Pilus, Stg fimbriae) (10, 60, 61) and several other adhesion factors involved in increased host colonization (including curli fimbriae, F9 fimbriae) (62, 63). Further experimental work should focus more on the key genetic determinants associated with highly abundant or prevalent taxa that allow for increased colonization across multiple hosts or environments, and assess how they are positioned within the broader metabolic and ecological network of the bovine intestine.

The differences between our two approaches may reflect the combined effects of sequencing depth limitations for shotgun metagenomics, or identification of closely related strains lacking the corresponding VFs. Nonetheless, besides our key-finding about important taxa and genes distributed differently across the two treatment groups, we also found that the overall virulence factor prevalence was substantially lower in ceftiofur-treated samples compared to control samples, and virulence factor prevalence in the antibiotic treatment group dropped after administration of ceftiofur. While this may seem beneficial for treating diseased animals, data from a similar pig experiment (64) indicate that antibiotic treatment can promote colonization by AMR *E. coli* that are strong colonizers. Since avirulent, AMR-carrying and pathogenic strains share intestinal niches and can horizontally exchange genes, antibiotic usage in farms may influence the emergence and evolution of multi-resistant pathogenic taxa. These dynamics warrant careful consideration in future experimental and surveillance studies.

## CONCLUSION

In this study, we assessed changes in the fecal microbiome of ruminating calves post ceftiofur antibiotic administration and *E. coli* administration. Our results show that ceftiofur administration is associated with distinct changes in microbial community composition, affecting key taxa as well as genes relevant to animal husbandry, with potential implications that may extend throughout the animal’s life and transmission to other hosts. However, despite the increased resolution of our methods, interpretation remains complicated mainly due to innate differences in the original microbiome communities likely stemming from environmental conditions, host microbiome structure and host genetics, and strain genetic background and other factors. Further research should focus on understanding the genetic determinants driving strain persistence, which will have broad implications for the control of multiresistant pathogenic taxa including *E. coli* in agricultural systems.

## DATA AVAILABILITY

Raw sequencing reads are available in the NCBI Sequence Read Archive (https://www.ncbi.nlm.nih.gov/sra) under BioProject accession numbers PRJNA1372467 (metagenomic reads) and PRJEB102994 (SCS reads). Scripts used for analysis and figure generation are available at https://forge.inrae.fr/sapountzis0454medis/hector-metagenomes.git.

## ETHICAL APPROVAL

Animal inoculation experiments were conducted in accordance with protocols approved by the State Office for Agriculture, Food Safety and Fisheries of Mecklenburg-Western Pomerania, Rostock, Germany (reference no. 7221.3-1-103019/20-1).

## SUPPLEMENTAL MATERIAL

**Figure S1:** Beta-diversity PCoA plots based on Euclidean distances of CLR-transformed MAG abundance profiles for the GWAS (**A**) and NJ (**B**) administration groups, after removing MAGs with <10% read-mapping coverage. For each administration group, two panels are shown: the top colored by treatment (green=control; orange=ceftiofur) and the bottom shaded by administration day. When all MAGs were included without the coverage filter, separation in the NJ group became less distinct, whereas patterns in the GWAS group remained largely unchanged (data not shown). **C**: PCoA of beta-diversity based on Euclidean distances of MAG abundance counts. Statistical tests and annotation are based on samples grouped by sampling timepoint (top) or animal ID number (bottom). Differences between groups were assessed using PERMANOVA, and group dispersion was evaluated using betadisper.

**Figure S2**: Identification of MAG prevalence across ceftiofur-exposed and control samples. (**A**). Heatmap showing the number of highly prevalent MAGs (>80% prevalence) unique to or shared between treatment groups. MAGs are grouped by family, or by the most specific available taxonomic level if a family-level classification was not possible. (**B**) Prevalence distribution of MAGs from each category (shared, highly prevalent in ceftiofur-exposed, highly prevalent in control) across the 25 control samples. (**C**) Prevalence distribution of MAGs from each category (shared, highly prevalent in ceftiofur-exposed, highly prevalent in control) across the 25 ceftiofur-exposed samples.

**Figure S3**: Distribution of MAGs encoding ARGs with at least 90% reference gene coverage. Heatmap shows the taxonomic identification of each MAG at the family level, when available. The numbers inside the boxes indicate the number of unique MAGs in which a given gene was identified.

**Figure S4**: Normalized abundances of CAG-485 (Muribaculaceae) MAGs encoding either the *cfxA5* (top MAG) or the *cfxA6* (bottom MAG) gene across each sampling date. The *cfxA5* (AY769934) gene was encoded with 94.2% identify by a MAG classified as CAG-485; the *cfxA6* (GQ342996) gene was encoded with 91.7% identify by a separate MAG classified as CAG-485 sp017547845.

**Figure S5:** Taxonomic Identification of all single-cells sequenced. Unidentified cells include any genome without either family or genus level identification. The number inside each box represents the count of cells classified as a given bacterial genus or family (if genus level identification is not available).

**Table S1**: Metadata for *E. coli* inoculated to calves in this study

**Table S2**: Metadata for all sequenced fecal samples in this study.

**Table S3: A:** Taxonomic summary and associated CheckM metrics of the 1607 MAGs with greater than 90% completeness and less than 5% contamination. **B**: MAGs identified as highly enriched in either control or ceftiofur-exposed samples via *Indicspecies* analysis (IndVal>0.80; fdr adjusted p<0.05). Statistical comparisons included only samples from 05d onwards.

**Table S4:** Taxonomic identification of sequenced single-cells through classification of genome k-mer signatures against the GTDB database.

**Supplementary File 1:** Mapping between genomes of administered *E. coli* genomes and denovo assembled contigs. Local BLAST Comparisons were used to identify: A) qualitatively and quantitatively the presence of the administered *E. coli* strains in our samples, B) chimeric contigs indicating potential Horizontal Gene Transfer events. Detailed descriptions of the methods and the results are provided, as well as two tables summarizing the results of A and B.

